# OSCAA: A Two-Dimensional Gaussian Mixture Model for Copy Number Variation Association Analysis

**DOI:** 10.1101/2023.09.25.559392

**Authors:** Xuanxuan Yu, Xizhi Luo, Guoshuai Cai, Feifei Xiao

## Abstract

Copy number variants (CNVs) are prevalent in the human genome which provide profound effect on genomic organization and human diseases. Discovering disease associated CNVs is critical for understanding the pathogenesis of diseases and aiding their diagnosis and treatment. However, traditional methods for assessing the association between CNVs and disease risks adopt a two-stage strategy conducting quantitative CNV measurements first and then testing for association, which may lead to biased association estimation and low statistical power, serving as a major barrier in routine genome wide assessment of such variation. In this article, we developed OSCAA, a flexible algorithm to discover disease associated CNVs for both quantitative and qualitative traits. OSCAA employs a two-dimensional Gaussian mixture model that is built upon the principal components from copy number intensities, accounting for technical biases in CNV detection while simultaneously testing for their effect on outcome traits. In OSCAA, CNVs are identified and their associations with disease risk are evaluated simultaneously in a single step, taking into account the uncertainty of CNV identification in the statistical model. Our simulations demonstrated that OSCAA outperformed the existing one-stage method and traditional two-stage methods by yielding a more accurate estimate of the CNV-disease association, especially for short CNVs or CNVs with weak signal. In conclusion, OSCAA is a powerful and flexible approach for CNV association testing with high sensitivity and specificity, which can be easily applied to different traits and clinical risk predictions.

## INTRODUCTION

Structure variants (SVs) are large-scale genetic variants that are increasingly recognized as contributors to human genetic variation and disease susceptibility as important as other genetic variants, such as single nucleotide polymorphisms (SNPs) [1]. Copy number variation (CNV), a type of SV that involves alteration in the DNA segments, has been extensively studied and shown to be associated with various human diseases. Common CNVs are of great significance in human genetic research, as they are more likely to influence genetic traits and disease risks than rare CNVs [2]. Studies have shown that common CNVs are associated with a wide range of human diseases, such as autism [3] schizophrenia [4] and cancers [5-7]. Furthermore, the discovery of common CNVs can aid in disease risk prediction, especially those that are potentially associated with the risk of certain diseases. Therefore, the identification and characterization of disease associated CNVs in human populations is crucial in advancing our understanding of human genetics and disease susceptibility, while accurate and robust methods for CNV detection and association testing are warranted for the development of effective disease risk prediction models and personalized medicine.

In the conventional methods of CNV-disease association testing, a two-stage strategy is often adopted, where CNVs are first detected, and their risks for diseases are then evaluated. For the first stage, various methods have been developed to detect CNVs for different genotyping platforms. For example, PennCNV [8] utilizes a hidden Markov model to detect CNVs for SNP array data, while DNAcopy [9] focuses on CNV detection for CGH array data using a circular binary segmentation (CBS) algorithm. ModSaRa [10] is an algorithm which was improved upon a local screening and ranking algorithm SaRa [11] to increase calling accuracy with optimized computational efficiency. ModSaRa can be applied to both SNP array and CGH array data and an improved version, modSaRa2 [12], was further developed to boost the power of detecting weak CNV signals by incorporating relative allele frequency. Additionally, CODEX [13] was designed for the whole exome sequencing (WES) data, adopting a Poisson latent factor model and CBS for normalization and chromosomal segmentation, respectively. Then, in the second stage, generalized linear models can be used to model the association between disease risk and copy number states in the shared CNV regions (CNVRs). However, the limitations of such two-stage CNV-disease association testing strategy are non-negligible, as misclassification of copy number states may occur during CNV calling, leading to wrongly assigned copy number states and subsequently biased association estimations if the incorrect results of CNV calling is not accounted for in the stage of association evaluation [14, 15].

There are existing methods accounting for uncertainties in CNV calling in the modelling of CNV-disease risk association, several of which adopted a one-stage framework. For example, CNVtest [16] utilized a scanning procedure to recursively select trait associated CNVs based on a score statistic without identifying sample specific CNVs. Similarly, without inferring the CNVs, Cheng et al. [17] employed a penalized linear regression model with two penalty terms to detect disease associated CNVs, in which the selection of CNVs shared by multiple samples and evaluation of the association with disease risks were achieved simultaneously. CNVtools [18] attempted to use the first principal component (PC) computed from probe intensities in a known CNVR, where a gaussian mixture model (GMM) was fitted to infer the latent copy number states, and the association with a disease was simultaneously tested by a likelihood ratio test. Although a linear discriminant function was used to further refine the first PC signal using the posterior probabilities of CNVs, misclassification of CNVs may still occur if the first PC is unable to capture the primary differences among CNVs. Our preliminary study suggested that even though the first two PCs were theoretically orthogonal, they were indeed correlated to some extent for samples with the same copy number state (Fig. S1A), mainly caused by the variation in mean signals across positions within the CNVR. Evidently, considering the correlation between the first two PCs enables better copy number profiling, which ultimately leads to improved robustness of the association testing (Fig. S1A, B).

In this study, we therefore develop the <u>O</u>ne-<u>S</u>tage <u>C</u>NV-disease <u>A</u>ssociation <u>A</u>nalysis (OSCAA) method, which utilizes a two-dimensional GMM model to assess the association between CNV and disease risk in a known CNVR with a one-stage framework. The two-dimensional GMM model is factorized into a signal model, a phenotype model, and a copy number model. To better differentiate samples with different copy numbers, we model the joint distribution of the first two PCs for samples with the same copy number in the signal model. The phenotype model comprises a linear model or a logistic model fitted for continuous or binary phenotype measurements, respectively. In addition, the phenotype model offers three hypotheses: the hypothesis of linear relationship between disease risk and copy number, the hypothesis of whether samples with abnormal copy number states are more/less likely to develop a disease, and the hypothesis of whether samples with deletions of copy numbers are more/less likely to develop a disease. The copy number model estimates the proportion of each copy number state and enables the evaluation of the potential impact of covariates on the proportions. Although we demonstrate the model based on SNP array data, our method can be extended to sequencing data with appropriate data normalization procedures.

The manuscript is organized as follows. The “Material and Methods” section introduces the OSCAA model, outlines the simulation designs, and provides a description of the melanoma microarray data [19]. In “Results” section, comprehensive simulations demonstrate the superiority of OSCAA over the existing one-stage method, CNVtools and two-stage methods, DNAcopy and CNVtest, both in CNV detection and association testing. Moreover, melanoma risk associated CNVs obtained from the analysis of melanoma data were consistent with prior findings. Finally, a brief conclusive remark is presented in the “Discussion” section.

## MATERIALS AND METHODS

### Two-dimensional Gaussian Mixture Model

For the *i*-th sample, let *x*_1*i*_, *x*_2*i*_ be the first two PCs derived from centered and standardized *log*_2_ ratio of probes within a specific CNVR, *y*_*i*_ be the phenotype indicating disease status, ***z***_***i***_ be the covariate vector of size *p*, and *s*_*i*_ indicates the unobserved copy number, where *i =* 1, …, *N* with *N* denoting the sample size. *s*_*i*_ can take on values of 0, 1, …, 4 representing deletion of double copies, deletion of single copy, normal/diploid, duplication of single copy, and duplication of double copies. Following the framework proposed by CNVtools [18], we jointly model *x*_1*i*_, *x*_2*i*_, *y*_*i*_, *s*_*i*_ *and* ***z***_***i***_ based on the following factorization:

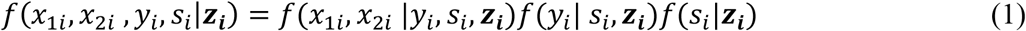

This model consists of three components, including 1) the signal model *f*(*x*_1*i*_, *x*_2*i*_ |*y*_*i*_, *s*_*i*_, ***z***_***i***_), 2) the phenotype model *f*(*y*_*i*_|*s*_*i*_, ***z***_***i***_), and 3) the copy number model *f*(*s*_*i*_| ***z***_***i***_).

The signal model *f*(*x*_1*i*_, *x*_2*i*_ |*y*_*i*_, *s*_*i*_, ***z***_***i***_) assumes that *x*_1*i*_ and *x*_2*i*_ follows a bivariate normal distribution with mean ***U*** _*i*_ and variance-covariance matrix **∑**_***i***_. The distribution of signals depends on the copy number but may also be influenced by phenotype *y*_*i*_ and extraneous covariates ***z***_***i***_, by accounting for which, we can adjust the difference among biological samples, or technical difference, etc. For simplicity, we demonstrate the signal model without including covariates ***z***_***i***_.

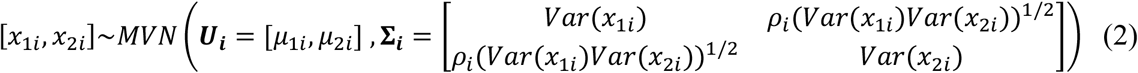

Specifically, the mean of each signal is modeled by a generalized linear model (GLM) with gaussian errors and an identity link function. Signal variance is estimated by a GLM with gamma errors and a logarithmic link function. In the covariance matrix, *ρ* measures the correlation between two PCs for samples with same copy number and is calculated based on the estimated means and variances. The signal mean models are separately expressed as:

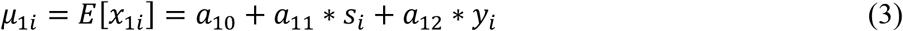

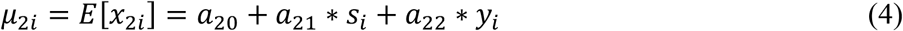

The signal variance models are:

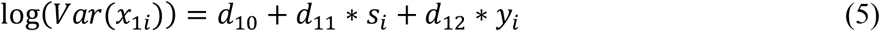

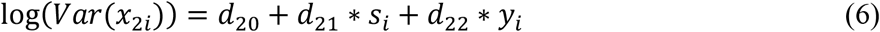

Where ***a***_**1**_ *=* (*a*_10_, *a*_11,_ *a*_12_), ***a***_**2**_ *=* (*a*_20_, *a*_21,_ *a*_22_) and ***d***_**1**_ *=* (*d*_10_, *d*_11,_ *d*_12_), ***d***_**2**_ *=* (*d*_20_, *d*_21,_ *d*_22_) are the mean and variance related parameters, respectively.

The phenotype model *f*(*y*_*i*_|*s*_*i*_, ***z***_***i***_) fits a logistic model for a binary phenotype or a linear model for a quantitative phenotype to model the association between phenotype and the latent copy number, adjusting for covariates ***z***:

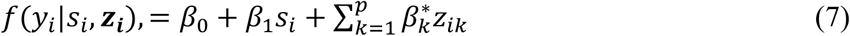

Where ***β*** *=* [*β*_0_, *β*_1_] *and* 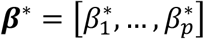 are regression coefficients. It is worth noting that ***s*** can be reparameterized so that different hypotheses can be tested. For instance, when *s*_*i*_ *=* 0 for copy number being 2 and *s*_*i*_ *=* 1 otherwise, the model tests whether there is a difference in the likelihood of developing a disease between samples with or without a CNV.

The copy number model *f*(*s*_*i*_| ***z***_***i***_) assumes that *s*_*i*_ follows a multinomial distribution. The proportions of copy numbers are allowed to be varied at different levels of ***z***_***i***_.

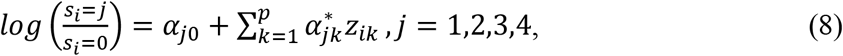

where ***α*** are regression coefficients.

### Model fitting and association testing

Note that the latent copy number *s* is unobserved during model fitting. We estimate the parameters using the maximum likelihood estimates (MLEs) via expectation conditional maximization (ECM) algorithm [20]. In the E step, posterior probabilities of each latent copy number are calculated for each sample based on current estimated parameters. In the CM step, three CM sub-steps are sequentially carried out with respect to the three factorized models. Within each CM sub-step, the parameters are estimated while keeping other parameters fixed. Once the algorithm converges, the Wald 𝒳^2^ test statistic is used to test the association between the disease and copy number, given the estimated 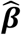 and variance-covariance matrix (Details in supplementary method).

### Correlation between decomposed components of genetic intensities from array data

We explored two array datasets to unveil the correlations between the decomposed components from the intensity data in a CNVR. We first investigated a CNVR consisting of 33 probes identified from the Wellcome Trust Case Control Study (WTCCC) [21] which contained 2,593 subjects. Additionally, in a genome-wide melanoma dataset that we have been frequently studying [19], in which CNVs were accurately detected using our previously developed method, modSaRa2 [12]. CNVRs were constructed using the RO method implemented in CNVRuler [22] and three CNVRs (chr2: 203607797-203618451, chr6: 77487750-77509808, chr9: 95540187-95541151) were randomly selected for our exploration in this study. For each of the identified CNVRs, CNVtools [18] was used to identify CNV state. The PCs were computed and further visualized in a two-dimensional plot and a histogram generated only using the first PC.

### Evaluation of OSCAA with existing one-stage method via simulation studies

We first evaluated the performance of OSCAA compared to the one-stage method of CNVtools. Following the spirit of that in CNVtools [18], without simulating the intensities for the whole sequence, we directly generated the signals of PCs defined in a CNVR for samples with different copy number states including deletion of one copy, normal/diploid, and duplication of one copy. The means and variances of the PCs for samples with normal states were estimated from the WTCCC dataset [21]. Additionally, the differences in means between adjacent copy number states was set as 1 for the first PC. In alignment with the reference CNVR [21], the difference in means was set as 0 for the second PC, while the correlation coefficient (*ρ*) was introduced to regulate the correlation between PCs. Based on the first PC, the signal-to-noise ratio (*Q*) measures the strength of noise for CNVs which is defined as the ratio between difference in means and the standard deviation. We evaluated the effect of varied values of *Q*, and *ρ* on the performance of OSCAA and CNVtools. Each simulation scenario was repeated 100 times and the mean ratio (*γ*) between estimated and true 𝒳^2^ test statistic was then calculated as a measurement of the consistency between the estimated and actual association testing results. A value of *γ* close to one indicates that the association has been estimated consistently. Furthermore, we employed receiver operating characteristic (ROC) curve and area under the curve (AUC) to compare the performance of the two methods.

To make fair comparison with CNVtools, signals of PCs for 2,000 samples were simulated for three different copy number states: deletion of one copy, normal/diploid, and duplication of one copy with proportion 50%, 30%, and 20%, respectively. The signal-to-noise ratio (*Q*) was generated for the first PC which ranged from 1.5 to 7.5. The values of PCs were simulated from bivariate normal distributions with mean vectors of (-0.4, 0), (0.6, 0), (1.6, 0) for deletion of one copy, normal state, and duplication of one copy, respectively. Additionally, the variance-covariance matrix used in the simulation was determined based on *Q* and *ρ*. Note that the correlation coefficient *ρ* was set to 0.8 when there was a correlation between the PCs. We then simulated a binary disease status for samples with or without CNV from Bernoulli distributions with varied parameter *p*, derived from the simplest logistic regression without the intercept term.

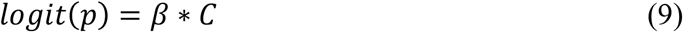

For the scenarios of alternative hypothesis that there was a CNV-disease association, we set odds ratio (OR) = *exp* (*β*) = 1.2. Moreover, we defined *C* = 1 for samples with a CNV and *C* = 0 for those without a CNV.

### Evaluation of OSCAA with existing two-stage methods via simulation studies

To further compared the performance of CNV detection and association testing between OSCAA and existing two two-stage methods, DNAcopy [9] and CNVtest [16], we explored the effect of varied CNV length, copy numbers, proportion of samples with the CNV, and combinations of different copy number states on the performance of these methods. Sensitivity, specificity, as well as ROC and AUC were evaluated for each method.

With 900 samples, CNV-free background signals of 6000 probes were first generated from a normal distribution with a mean of 0 and a standard deviation (SD) of 0.18. We then assigned a binary disease status for these samples of cases and controls with equal sizes. Next, 120 CNVRs were chosen from 6000 probes with different CNV lengths (short: 5∼20 probes, medium: 20∼50 probes and long: 50∼80 probes). For each of the 120 CNVRs, we assigned CNV status for samples using a binary distribution with *p* derived from the logistic model, as shown in Eq.9 with varied CNV proportions (5%, 10%, 20%, 30%, 50%) and ORs (1.5 for strong association, 1.2 for weak association, and 1 for no association). Finally, signals of CNVs were added to the background signal, varying the copy number states with means and SDs provided by the Illumina website ([23], Table S1). The copy number states included deletion of double copies (Del.D), deletion of single copy (Del.S), duplication of single copy (Dup.S) and duplication of double copies (Dup.D). In addition, we also assigned mixed copy number states to samples per CNVR, including a mixture of two deletion states (Del.D and Del.S), a mixture of two duplication states (Dup.S and Dup.D), and a mixture of four copy number states (Del.D, Del.S, Dup.S and Dup.D).

### Application to the whole genome melanoma dataset

We applied OSCAA to a whole genome data from Gene Environmental Association Studies initiative (GENEVA), which included 3,115 participants. High-density SNP array data were obtained using the Illumina Omni1-Quad_v1-0_B array platform for patients with skin cutaneous melanoma and healthy controls. The details of this dataset have been previously described in Amos et al. [19]. We applied the same quality control procedures as described in modSaRa2 [12], resulting in a total of 2,829 sample with 1,109,421 markers in each sample.

In total, 354,210 CNVs were identified and CNVRs shared among samples were constructed by utilizing the reciprocal overlap (RO) method implemented in the CNVRuler software [22]. The RO method defines the CNVR based on the degree of overlapping between any two contributing CNVs. We used a default threshold of the reciprocal overlapping region of 50%.

For each of the defined CNVRs, we applied OSCAA to evaluate the association between CNVs and melanoma risk. Two models were used: (1) Model 1 in which deletions and duplications were assumed to have the same effect on the risk of melanoma; (2) Model 2 in which only deletions were associated with the risk of melanoma. In each model, gender was adjusted in the signal model, whereas gender and age were included as covariates in the phenotype model.

To ensure the reliability of association testing results, we further conducted a quality control procedure to filter out CNVRs of poor quality. These CNVRs exhibited a characteristic where only part of the region harbored the CNV signal. Consequently, more noise was introduced, making it difficult for the PCs to effectively distinguish between these samples and ultimately leading to biased association estimation. To resolve this problem, we introduced the F score, which measured the clustering quality for a CNVR, with a higher value indicating better clustering quality. Explicitly, the F score was calculated as the ratio of mean square errors (MSE) comparing between- and within-cluster MSE using the mean of the CNV signal. The threshold for F score was empirically determined by visualizing raw signals for CNVRs so that CNVRs with good clustering quality could be retained. In this study, the threshold for F score was set as 3,000.

## RESULTS

### Correlation exists between decomposed components of genetic intensities from array data

The motivation of our methodology was inspired by the discovery of the correlated structure within the decomposed components obtained from the real genetic intensity data. As discussed, in the existing one stage association method CNVtools [18], the CNV states were identified only based on the first PC (Fig. S1A, C). However, when visualizing the identified CNV states in a two-dimensional plot using the first two PCs, it was observed that a considerable proportion of samples with normal and duplication states were not adequately distinguished from each other. Furthermore, evident correlations were observed between two PCs for samples with same copy number states (Fig. S1B). The clear boundary observed between clusters suggested that by utilizing two PCs while considering the correlations, samples with varied copy number states would be effectively distinguished from each other (Fig. S1B). This pattern was also extensively observed for CNVs in the melanoma dataset, such as those in chr2: 203607797-203618451 (Fig. S1C-D), chr6: 77487750-77509808 (Fig. S1E-F) and chr9: 95540187-95541151 (Fig. S1G-H). In addition to the correlations observed for the CNVs (chr6: 77487750-77509808, chr9: 95540187-95541151), the method utilizing only one PC exhibited a tendency to select a larger number of clusters. Consequently, individuals with the same CNV state were assigned to different clusters (Fig. S1E-H). Rather, by employing two PCs for CNV detection, samples in the WTCCC dataset were effectively distinguished, with the Pearson’s correlation coefficients being 0.62, 0.67 and 0.63 for samples with deletions, normal and duplications, respectively. Similarly, in the melanoma data (chr2: 203607797-203618451), the Pearson’s correlation coefficients between PCs were 0.8, 0.8, and 0.04 for samples with normal state, single deletions, and double deletions. Moreover, three clusters were identified for the two CNVs (chr6: 77487750-77509808, chr9: 95540187-95541151), aligned with the observed patterns. Together, these findings provided strong evidence of the existence of correlation among samples with the same copy numbers.

### Simulations showed superior performance of OSCAA in comparison to one-stage method

Overall, OSCAA outperformed CNVtools in scenarios where there was a correlation between two PCs and the signal-to-noise ratio (Q) was small (Fig. 1). Specifically, OSCAA maintained its power of detecting CNV-disease risk associations, while CNVtools tended to underestimate the test statistic, leading to significant loss of power (Fig. 1E). For example, when Q = 1.5, OSCAA yielded a *γ* of 1.00, while CNVtools had a *γ* of 0.76. Besides, in cases where there was no CNV-disease association, both methods overestimated the test statistic for small *Q* values. However, OSCAA still showed better controlled type I error compared to CNVtools (Fig. 1F). For instance, when Q was 1.5, OSCAA had a *γ* of 3.60, while CNVtools had a *γ* of 21.84. With large Q values, where samples with different copy numbers could be well distinguished based solely on the first PC, both methods yielded consistent estimations (Fig. 1B, D).

We also evaluated the performance of OSCAA under the condition of no correlation among samples with the same copy number states. OSCAA and CNVtools showed comparable performance in the association testing power. Specifically, CNVtools tended to underestimate the test statistic when there was an association, while OSCAA showed an overestimation but more consistent results across different signal-to-noise ratios (Fig. S2E). When there was no association, both methods presented overestimation of test statistics, resulting in an inflation of type I error for small signal-to-noise-ratio (Fig. S2F). The ROC curves for selected scenarios further highlighted the superiority of OSCAA over CNVtools in CNV-disease risk association testing, especially for small Q values and correlated PCs (Fig. 2).

In summary, OSCAA presented significantly better performance than CNVtools when there was a correlation between two PCs among samples with the same copy number, while maintaining comparable performance when no correlation existed between the two PCs.

### Simulations showed superior performance of OSCAA in comparison to two-stage methods

For detecting strong CNV-disease associations, OSCAA demonstrated superior performance than DNAcopy and CNVtest particularly for short CNVs with weak signals (i.e., single duplication) (Fig. 3A). Also, superior performance of OSCAA was observed when a certain proportion of CNVs were with weak signals (Fig. 3A, D, F). For example, when the CNV consisted of only a single copy of duplications (Fig. 3A), OSCAA presented better performance in detecting disease associated CNVs. Across all CNV proportion settings, OSCAA consistently achieved the highest AUCs outperforming DNAcopy and CNVtest by more than 10% in AUC when CNV proportion increased to 0.5. Such difference was also highlighted in the ROC curves (Fig. 4). For CNVs with strong signals, such as deletion of single copy (Fig. 3B), duplications/deletions of double copies (Fig. 3C), and a mixture of these signals (Fig. 3E), all three methods yielded comparable performance. Furthermore, three methods showed similar performance when the CNV length was medium or long (Fig. S3).

Consistently, when detecting the effect from relatively weak CNV-disease associations, OSCAA still outperformed DNAcopy and CNVtest for short CNVs with weak signals or when a certain proportion of CNVs were weak signals (Fig. S4A, D, F). Simulations also evaluated these methods in association testing of medium and long CNVs, where CNVs presented three different levels of weak mean signals (Fig. S6). The results revealed that OSCAA consistently outperformed the other two methods in these scenarios. However, for short CNVs with very weak signal (Fig. S6A), all three methods failed to accurately detect CNVs and furthermore their associations. It is also noted that as CNV length increased, the performance of these methods converged to similar levels.

In brief, by incorporating the uncertainty in CNV calling into the modelling of CNV-disease association, OSCAA demonstrated advantageous performance over DNAcopy and CNVtest in assessing associations for CNVs with weak signals.

### Analysis of whole genome melanoma dataset

In total, 22,342 CNVRs were constructed in autosomes for 2,829 samples using the RO method from CNVRuler [22]. Although CNVRs overlapped with both protein coding and non-coding regions, overlapping protein coding regions contributed more to the variation in phenotypes than the non-protein regions [12]. Therefore, the CNVRs were mapped to the coding genes using NCBI build 36 (hg18) reference. Overall, after conducting the quality control procedure, 54 statistically significant genes (*P*-value < 0.05) were retained in the duplication/deletion equal effect model (Model 1) (Table 1, Table S2) and 26 were retained in the deletion-only effect model (Model 2) (Table S3). Among the top 20 significant genes from the two models, only three genes (*C1ORF159, ARID1B, TECPR1*) overlapped, suggesting that duplications did not have a risk effect on melanoma development. Clinical associations between the detected genes and melanoma risk have been reported in previous studies, most of which used single nucleotide polymorphism data. For example, deletion of regions in 1p36.3 was shown to be a frequent event in malignant melanoma [24] and uveal melanoma [25], which harbors *MXRA8* detected by Model 1 (OR = 1.47, 95% CI = 1.24-1.76, *P*-value = 1.54×10^−5^), *TTLL10* (OR = 2.17, 95% CI = 1.29-3.65, *P*-value = 3.58×10^−3^), *TNFRSF18* (OR = 1.73, 95% CI = 1.09-2.75, *P*-value = 2.06×10^−2^) detected by Model 2, and *C1ORF159* (OR = 2.54, 95% CI = 1.45-4.47, *P*-value = 1.18×10^−3^) detected by both models. Additionally, deletion in 10q26.3 (OR = 1.27, 95% CI = 1.08-1.51, *P*-value = 4.5×10^−3^), which harbored *PWWP2B*, was also detected in both models and shown to be related to metastasis melanoma [26].

In conclusion, the application of OSCAA in a real melanoma case control study demonstrated the effectiveness of OSCAA in detecting disease associated CNVs, though further evaluation was still needed for a solid conclusion for the scientific discoveries. In addition, the quality control procedure developed in our study was found to be an efficient step in filtering out low-quality CNVRs.

## DISCUSSION

In this paper, we focus on developing a one-stage strategy that offers a robust method for simultaneous CNV detection and CNV-disease association testing by fitting a two-dimensional GMM model which considers the first two PCs and their correlations for a CNVR. In contrast to traditional association methods that use a two-stage framework involving CNV detection and subsequent regression modeling, our approach considers the probabilities of false negative or false positive calls, leading to more accurate CNV calls and more reliable association testing results.

Our comprehensive simulation studies demonstrated the remarkable performance of OSCAA in comparison to the only existing one-stage method, CNVtools. OSCAA exhibited a significant increase in statistical power with effectively controlled type I error, particularly when there was a correlation between PCs and a small signal-to-noise ratio. Furthermore, OSCAA outperformed existing two-stage methods, CNVtest and DNAcopy, in both CNV detection and association testing, especially for CNVs characterized by weak signals and short lengths. To further validate OSCAA’s capabilities, we applied it to a whole-genome microarray dataset of cutaneous melanoma and obtained consistent results of disease-related CNVs with previous studies. These results solidify OSCAA’s superiority and robustness across various scenarios. OSCAA can be applied to data generated by multiple microarray platforms that follow a Gaussian distribution and can also be extended to sequencing datasets with appropriate data transformations.

OSCAA does have certain limitations. Firstly, it relies on the availability of a predefined CNVR, which may not always be available or well-defined. Therefore, efficient CNVR construction methods are still needed. Moreover, despite the advantage of incorporating more PCs and considering the correlations between PCs in the model, there remains a possibility of information loss if a considerable proportion of total variance remains uncaptured by the first two PCs. However, caution is needed when incorporating additional PCs as that may lead to increased computational burden and potential risk of noise enlargement.

## ACKNOWLEDGEMENTS

We acknowledge Gene Environment Association Studies initiative (GENEVA) for providing the melanoma dataset for our study.

## CODE AVAILABILITY

The source code for OSCAA can be accessed from https://github.com/FeifeiXiao-lab/OSCAA.

