## Supplementary material for "OSCAA: A Two-Dimensional Gaussian Mixture Model for Copy Number Variation Association Analysis"

### Materials and methods

#### Model fitting and association testing

As the latent copy number $s$ is unobserved "missing data", the complete data log likelihood is defined as

$$\sum_{i=1}^{N} log\left(\sum_{s_i=0}^{4} f(x_{1i}, x_{2i}, y_i, s_i|\boldsymbol{z_i}, \boldsymbol{a_1}, \boldsymbol{a_2}, \boldsymbol{d_1}, \boldsymbol{d_2}, \boldsymbol{\rho}, \boldsymbol{\alpha}, \boldsymbol{\beta})\right)$$

$$= \sum_{i=1}^{N} log\left(\sum_{s_i=0}^{4} f(x_{1i}, x_{2i} | y_i, s_i, \boldsymbol{z_i}) f(y_i| s_i, \boldsymbol{z_i}) f(s_i|\boldsymbol{z_i})\right) \quad (1)$$

Parameters in the model are estimated using the maximum likelihood estimates (MLEs) via expectation conditional maximization (ECM) algorithm [1]. In the E step, based on current estimated parameters, posterior probabilities of each latent copy number are calculated for each sample. In the CM step, three CM steps are carried out with respect to the three factorized models, sequentially. Within each CM step, the parameters are estimated given other parameters fixed.

Explicitly, in the E step, the posterior probability for copy number $s_i$ in sample $i$ is computed as

$$P_{is} = \frac{p(x_{1i}, x_{2i}, y_i, s_i|\boldsymbol{z_i}, \boldsymbol{a_1}, \boldsymbol{a_2}, \boldsymbol{d_1}, \boldsymbol{d_2}, \boldsymbol{\rho}, \boldsymbol{\alpha}, \boldsymbol{\beta})}{\sum_{s_i=0}^{4} p(x_{1i}, x_{2i}, y_i, s_i|\boldsymbol{z_i}, \boldsymbol{a_1}, \boldsymbol{a_2}, \boldsymbol{d_1}, \boldsymbol{d_2}, \boldsymbol{\rho}, \boldsymbol{\alpha}, \boldsymbol{\beta})} \quad (2)$$

In the CM steps, the parameters in each CM step are estimated by maximizing the expectation of "complete" data log likelihood in turn.

$$\sum_{i=1}^{N} \sum_{s_i=0}^{4} P_{is} log\left(p(x_{1i}, x_{2i}, y_i, s_i|\boldsymbol{z_i}, \boldsymbol{a_1}, \boldsymbol{a_2}, \boldsymbol{d_1}, \boldsymbol{d_2}, \boldsymbol{\rho}, \boldsymbol{\alpha}, \boldsymbol{\beta})\right) \quad (3)$$

To fit the signal mean models, we maximize the expected log likelihood with respect to $\boldsymbol{a_1}$ and $\boldsymbol{a_2}$, respectively. For example, for modeling the first PC, $\boldsymbol{a_1}$ can be estimated by maximizing the objective function $Q(\boldsymbol{a_1})$.

$$Q(\boldsymbol{a_1}) = \sum_{i=1}^{N} \sum_{s=0}^{4} P_{is} \left( \left( \frac{x_i - a_{10} - a_{11} * s_i - a_{12} * y_i}{exp\left(\frac{1}{2}(d_{10} + d_{11} * s_i + d_{12} * y_i)\right)} \right)^2 \right)$$

$$= \sum_{i=1}^{N} \sum_{s=0}^{4} \frac{P_{is}}{exp(d_{10} + d_{11} * s_i + d_{12} * y_i)} (x_i - a_{10} - a_{11} * s_i - a_{12} * y_i)^2 \quad (4)$$

Obviously, $\boldsymbol{a_1}$ can be estimated by fitting a weighted GLM for $x_{1i}$ with gaussian errors and an identity link function, where the weight is $\frac{P_{is}}{Var(x_{i1})}$. Likewise, $\boldsymbol{a_2}$ is estimated by utilizing the same approach.

To fit the signal variance models, maximize the expected log likelihood with respect to $\boldsymbol{d_1}$, and $\boldsymbol{d_2}$, respectively. For parameters in the signal variance model for the first PC, they are estimated by maximizing the objective function $Q(\boldsymbol{d_1})$.

$$Q(\boldsymbol{d_1}) = \sum_{i=1}^{N} \sum_{s=0}^{4} \frac{P_{is}}{\exp\,(d_{10} + d_{11} * s_i + d_{12} * y_i)} \exp\left( \frac{(x_{1i} - \mu_{1i})^2}{\exp(d_{10} + d_{11} * s_i + d_{12} * y_i)} \right) \quad (5)$$

Note that $\frac{1}{\exp\,(d_{10} + d_{11} * s_i + d_{12} * y_i)} \exp\left( \frac{(x_{1i} - \mu_{1i})^2}{\exp(d_{10} + d_{11} * s_i + d_{12} * y_i)} \right)$ is the kernel density of $Gamma(\exp(d_{10} + d_{11} * s_i + d_{12} * y_i), 1)$ in the form of shape-scale parametrization with the random variable being $(x_{1i} - \mu_{1i})^2$, shape parameter $v = 1$, and scale parameter $u_i = \exp(d_{10} + d_{11} * s_i + d_{12} * y_i)$. Therefore, $\boldsymbol{d_1}$ or $\boldsymbol{d_2}$ can be estimated by fitting a weighted GLM for $(x_{1i} - \mu_{1i}))^2$ or $(x_{2i} - \mu_{2i}))^2$ with gamma errors and weight, $P_{is}$, respectively. Finally, $\rho_i$ is estimated nonparametrically for each latent copy number using samples at the same covariates level.

To fit the phenotype model, a weighted logistic model for a binary phenotype or a weighted GLM for a quantitative phenotype will be fitted with weight being $P_{is}$.

To fit the copy number model, a multinomial logistic model will be fitted with weight, $P_{is}$. Then the probability of $s_i$ for each sample is calculated by using the Softmax function. For example, set $s_i = 0$ as the reference level,

$$\begin{cases} p(s_i = 0) = \frac{1}{1+\sum_{j=1}^{4} \exp\,(\alpha_{j0}+\sum_{k=1}^{p} \alpha_{jk}^{*} z_{ik})}, if\ s_i = 0 \\ p(s_i = j) = \frac{\exp\,(\alpha_{j0}+\sum_{k=1}^{p} \alpha_{jk}^{*} z_{ik})}{1+\sum_{j=1}^{4} \exp\,(\alpha_{j0}+\sum_{k=1}^{p} \alpha_{jk}^{*} z_{ik})}, otherwise \end{cases} \quad (6)$$

To test the significance of the association between phenotype and copy number state $H_0: \beta = 0\ vs\ H_1: \beta \neq 0$, a Wald $\chi^2$ test statistic and the corresponding $p$-value are computed given the estimated $\hat{\beta}$ and variance-covariance matrix.

#### Supplementary Figures and Tables

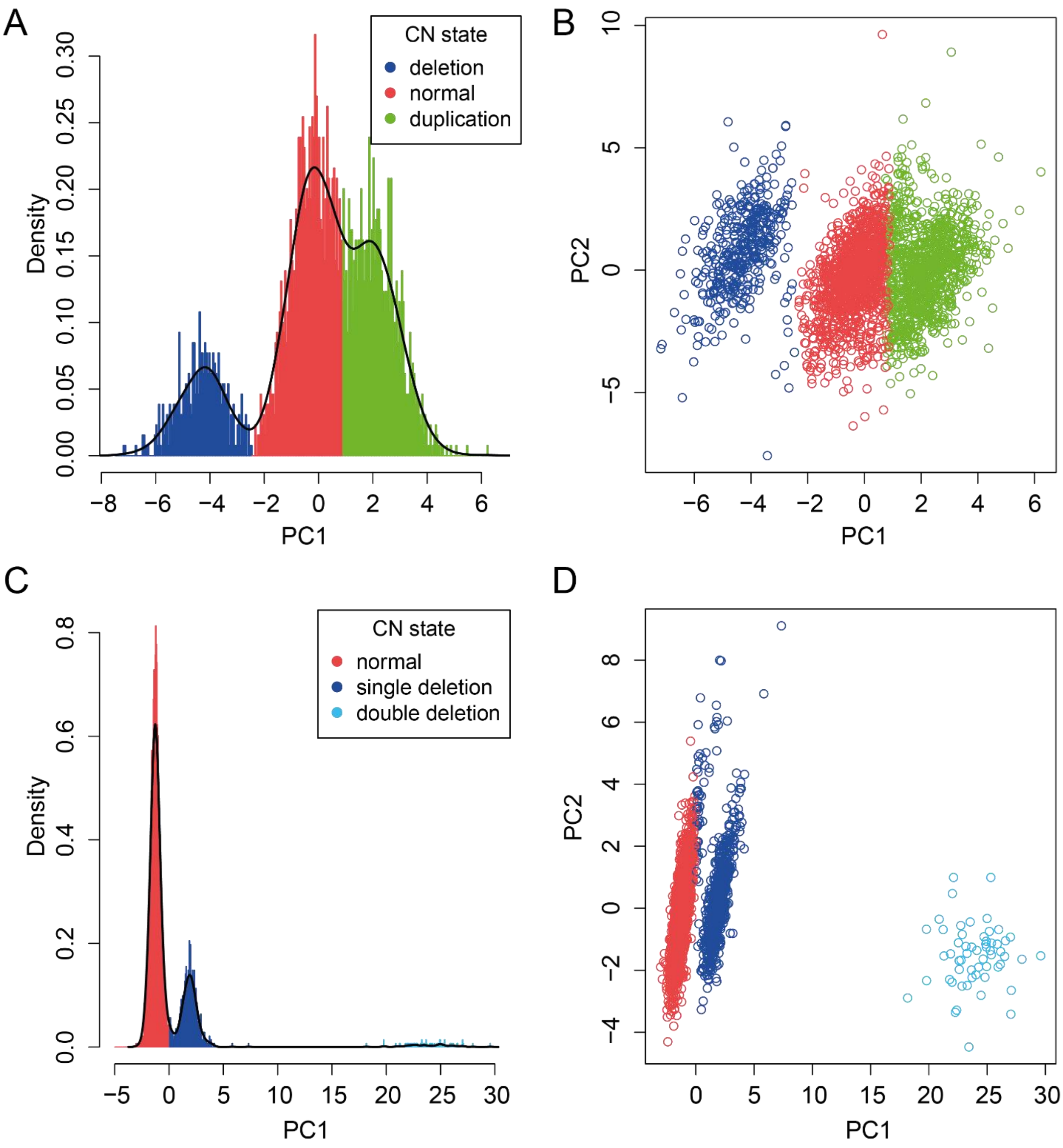

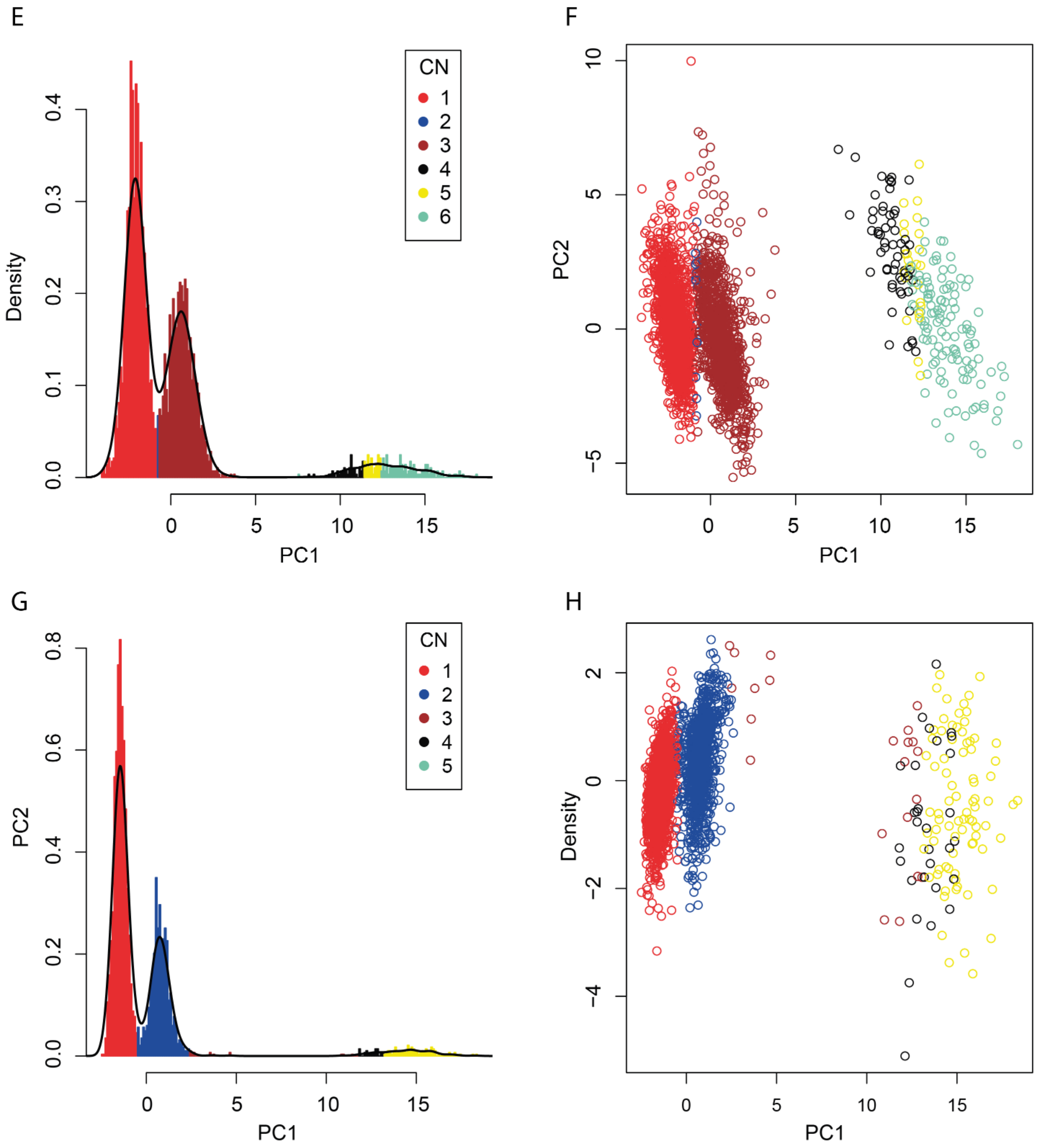


**Supplementary Fig. S1. Principal component analysis for A112 CNV in WTCCC data and a CNV in melanoma data.** The first two PCs were computed for the A112 CNV consisting of 33 probes in WTCCC data (A, B), and three identified CNV in melanoma data (C-H). CN states were identified by using the first PC using CNVtools, PC: principal components.

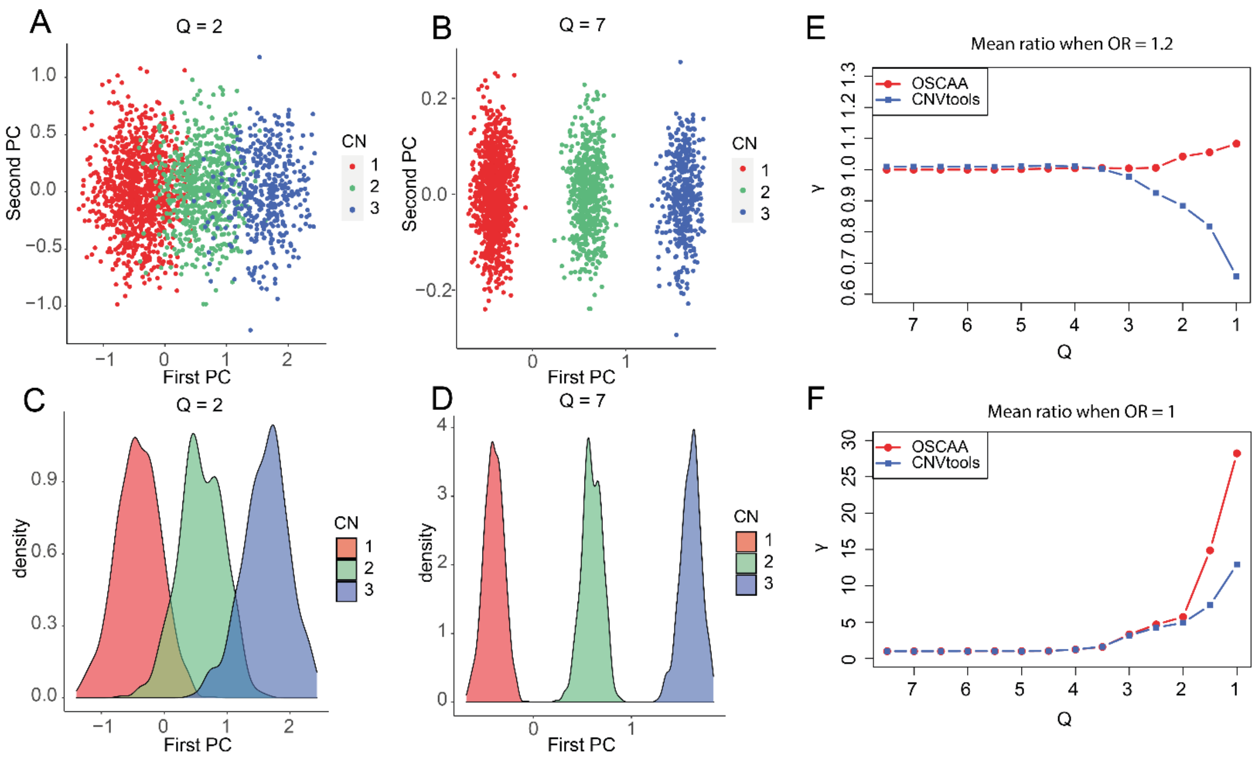


**Supplementary Fig. S2. Performance evaluation of OSCAA with CNVtools via simulation studies.** Assume there was no correlation between the first and second PCs. Signals for 2,000 samples were simulated for 3 different CNV states: deletions, normal, and duplications from bivariate normal distributions with parameters derived based on $Q$. The CNV proportion varied among 50%, 30%, and 20%. $Q$ denoted the signal-to-noise ratio ranging from 1 to 7.5. A binary disease status was simulated; OSCAA was fitted using the first two PCs while CNVtools was fitted by only using the first PC. Distribution of CNV signals were displayed for $Q$ = 2 (A, C) and 7 (B, D), respectively. Mean ratio ($\gamma$) was used to evaluate the performance of each method under various $Q$ values when OR = 1.2 (E) and OR = 1 (F). $\gamma$: mean ratio between estimated $\chi^2$ and true $\chi^2$ across 100 simulations. OR: Odds Ratio.

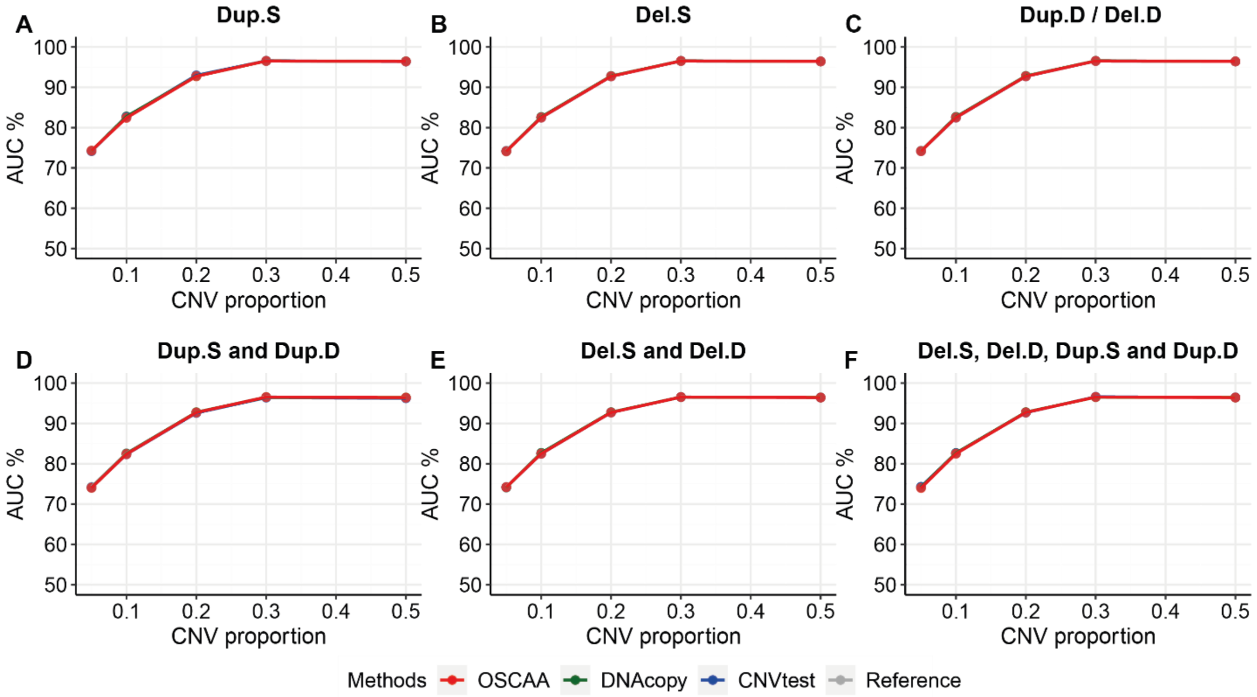


**Supplementary Fig. S3. Performance evaluation of OSCAA with two-stage methods in medium/long CNVs via simulation studies with $OR_{h1}$ = 1.5.** Signals of 120 CNVRs of medium CNV length (20~50 probes) or long CNV length (50~80 probes) were simulated for varied copy number states including Dup.S (A), Del.S (B), Del.D/ Dup.D (C), mixture of 2 duplication states (D: Dup.S and Dup.D), mixture of 2 deletion states (E: Del.D and Del.S); and mixture of 4 CNV states (F: Del.D, Del.S, Dup.S and Dup.D). The CNV proportion ranged from 0.05 to 0.5. In each scenario, a binary disease status was simulated for each sample with OR = 1.5 or OR = 1. The reference line (grey line) represented the actual AUCs with each CNV correctly detected. Del.D: deletion of double copies; Del.S: deletion of single copy; Dup.S: duplication of single copy; Dup.D: duplication of double copies; OR: Odds Ratio; SD: standard deviation; CNVR: CNV region; AUC: area under curve.

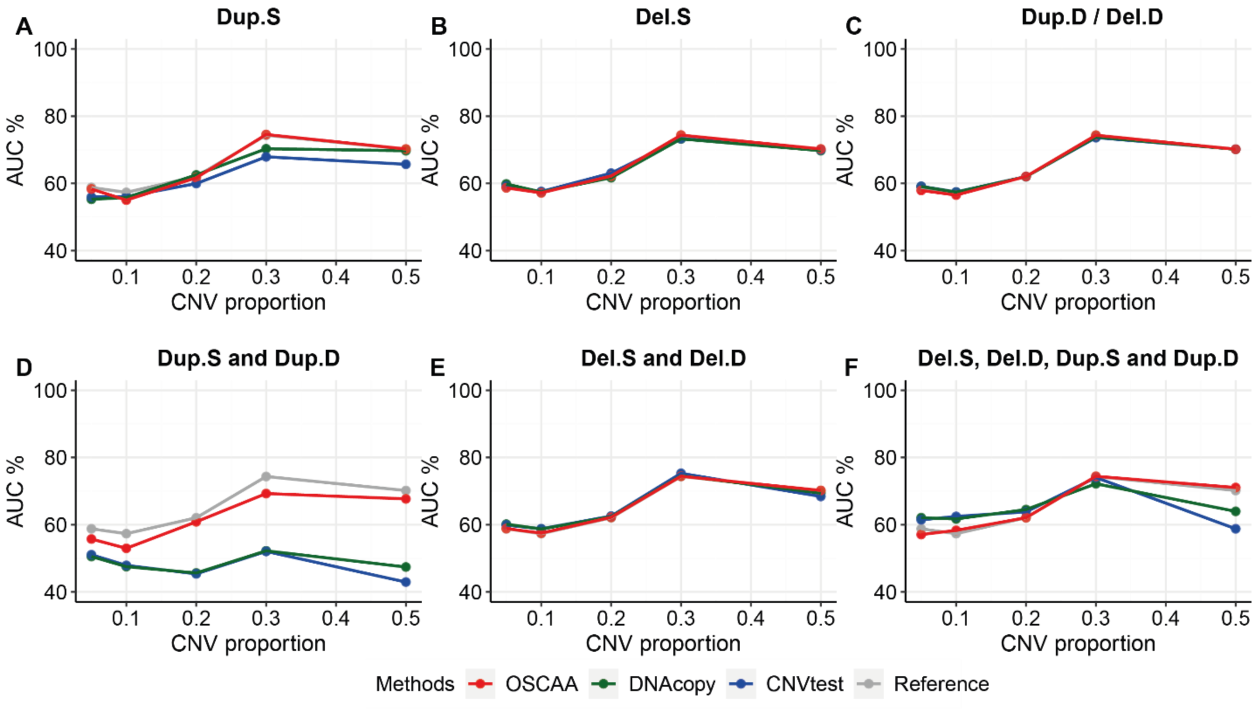


**Supplementary Fig. S4. Performance evaluation of OSCAA with two-stage methods, DNAcopy and CNVtest in short CNVs via simulation studies with $OR_{h1}$ = 1.2.** Signals of 120 CNVRs of short CNV length (5~20 probes) were simulated for varied copy number states including Dup.S (A), Del.S (B), Del.D/ Dup.D (C), mixture of 2 duplication states (D: Dup.S and Dup.D), mixture of 2 deletion states (E: Del.D and Del.S); and mixture of 4 CNV states (F: Del.D, Del.S, Dup.S and Dup.D). The CNV proportion ranged from 0.05 to 0.5. In each scenario, a binary disease status was simulated for each sample with OR = 1.5 or OR = 1. The reference line (grey line) represented the actual AUCs with each CNV correctly detected. Del.D: deletion of double copies; Del.S: deletion of single copy; Dup.S: duplication of single copy; Dup.D: duplication of double copies; OR: Odds Ratio; SD: standard deviation; CNVR: CNV region; AUC: area under curve.

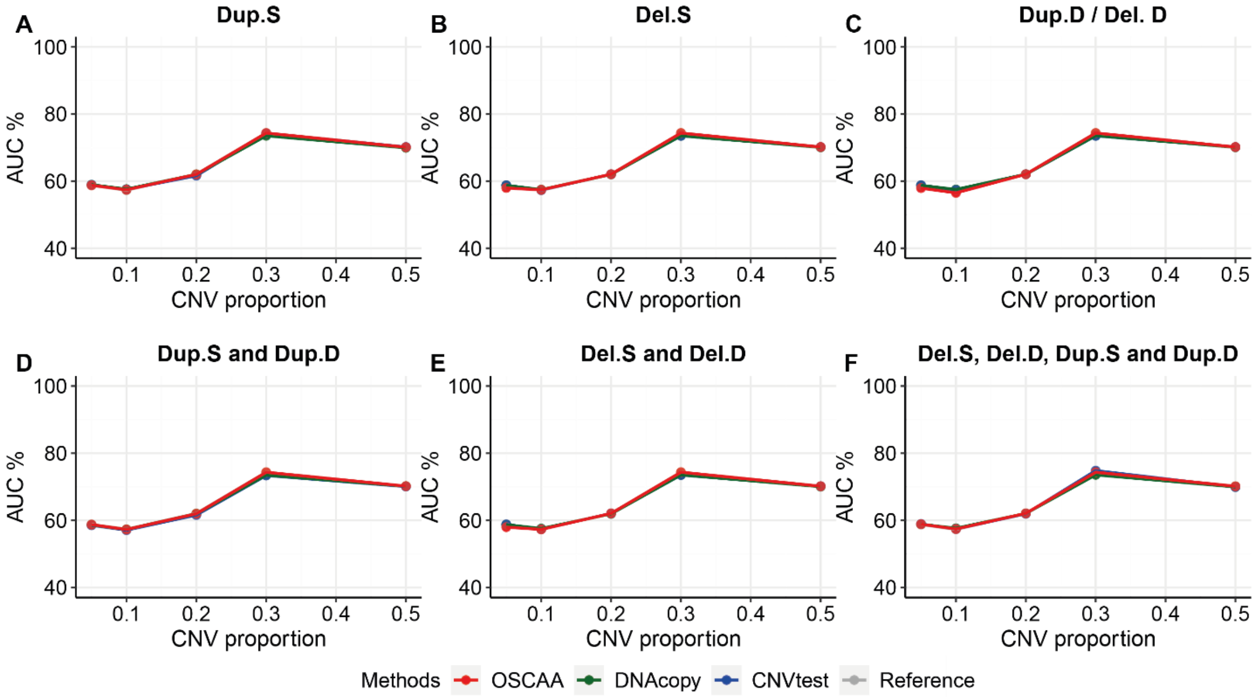


**Supplementary Fig. S5. Performance evaluation of OSCAA with two-stage methods, DNAcopy and CNVtest in medium/long CNVs via simulation studies with $OR_{h1}$ = 1.2.** Signals of 120 CNVRs of medium CNV length (20~50 probes) or long CNV length (50~80 probes) were simulated for varied copy number states including Dup.S (A), Del.S (B), Del.D/ Dup.D (C), mixture of 2 duplication states (D: Dup.S and Dup.D), mixture of 2 deletion states (E: Del.D and Del.S); and mixture of 4 CNV states (F: Del.D, Del.S, Dup.S and Dup.D). The CNV proportion ranged from 0.05 to 0.5. In each scenario, a binary disease status was simulated for each sample with OR = 1.2 or OR = 1. The reference line (grey line) represented the actual AUCs with each CNV correctly detected. Del.D: deletion of double copies; Del.S: deletion of single copy; Dup.S: duplication of single copy; Dup.D: duplication of double copies; OR: Odds Ratio; SD: standard deviation; CNVR: CNV region; AUC: area under curve.

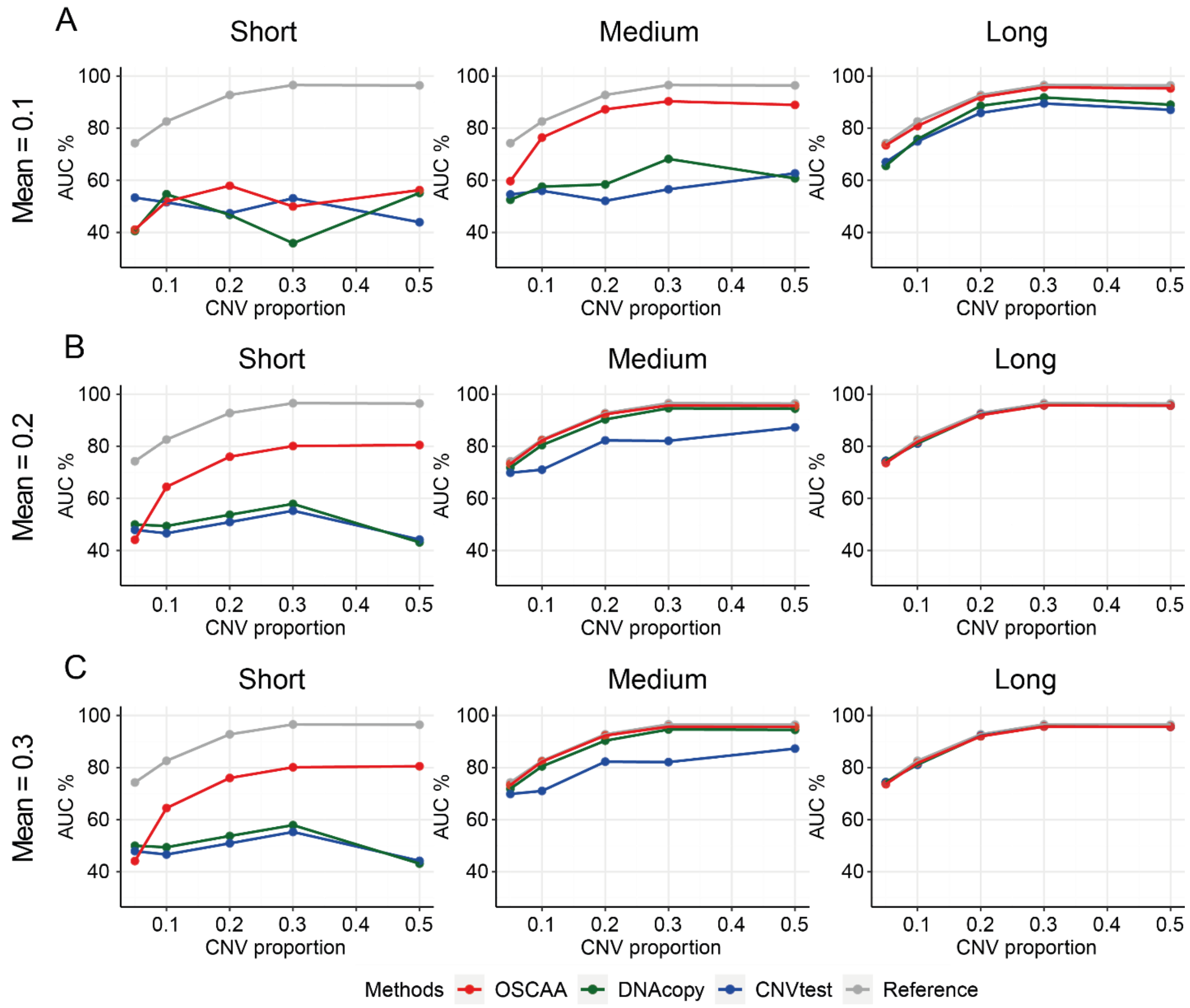


**Supplementary Fig. S6. Performance evaluation of OSCAA with two-stage methods, DNAcopy and CNVtest via simulation studies for CNVs with weak signals.** Signals of 120 CNVRs are added to the background CNV-free signal for n% (n=5, 10, 20, 30, 50) of 900 samples, with varied CNV length (short: 5~20 probes, medium: 20~50 probes, long: 50~80 probes) and varied levels of CNV signals including mean = 0.1 (A), 0.2 (B), and 0.3 (C). Then, a binary disease status was simulated for each sample from a binomial distribution with OR = 1.5 or OR = 1. Del.D: deletion of double copies; Del.S: deletion of single copy; Dup.S: duplication of single copy; Dup.D: duplication of double copies; OR: Odds Ratio; SD: standard deviation; CNVR: CNV region; AUC: area under curve.

**Supplementary Table S1. Copy number states with empirical parameter estimates**

| Copy number state | LRR mean | LRR SD |
|---|---|---|
| Del. D | -5 | 0.18 |
| Del. S | -0.45 | 0.18 |
| Dup. S | 0.3 | 0.18 |
| Dup. D | 0.75 | 0.18 |

LRR: log R ratio; SD: standard deviation; Del.D: deletion of double copies; Del.S: deletion of single copy; Dup.S: duplication of single copy; Dup.D: duplication of double copies;

**Supplementary Table S2. Significantly associated germline CNVRs with melanoma risk detected by Model 1.** CNV calls were first generated by using modSaRa, and CNVRs were then defined by using RO method by using CNVRuler. OSCAA was applied to each CNVR adjusting age and gender effect. NCBI build 36 (hg18) was used to find overlapped genes for CNVRs with significant CNV-disease association (p value < 0.05). 53 significant genes were shown in the table. RO: reciprocal overlap; CNVR: CNV region.

| Gene | Chr | Coordinates | OR (95% CI) | P value |
|---|---|---|---|---|
| MXRA8 | 1p36.33 | 1277934-1283778 | 1.47(1.24, 1.76) | 1.54E-05 |
| GPR35 | 2q37.3 | 241217468-241219342 | 2.72(1.57, 4.73) | 3.80E-04 |
| TXNDC2 | 18p11.22 | 9875723-9878156 | 0.45(0.29, 0.70) | 4.59E-04 |
| MACROD2 | 20p12.1 | 13924146-15981841 | 0.76(0.65, 0.89) | 5.66E-04 |
| C1ORF159 | 1p36.33 | 1007061-1041599 | 2.61(1.49, 4.56) | 7.71E-04 |
| ARFGEF1 | 8q13.2 | 68272451-68418466 | 0.63(0.48, 0.83) | 7.76E-04 |
| ARID1B | 6q25.3 | 157140778-157572094 | 2.39(1.41, 4.05) | 1.23E-03 |
| OR7C2 | 19p13.12 | 14913301-14914260 | 1.29(1.10, 1.50) | 1.29E-03 |
| TECPR1 | 7q21.3 | 97683983-97719404 | 1.26(1.08, 1.48) | 4.18E-03 |
| PWWP2B | 10q26.3 | 134060692-134081348 | 1.27(1.08, 1.51) | 4.50E-03 |
| OR4N2 | 14q11.2 | 19365448-19366371 | 2.52(1.32, 4.82) | 5.08E-03 |
| TCF15 | 20p13 | 532637-538910 | 2.12(1.25, 3.59) | 5.46E-03 |
| NCS1 | 9q34.11 | 131974678-132039404 | 1.28(1.08, 1.53) | 5.68E-03 |
| CDC34 | 19p13.3 | 482733-493087 | 2.01(1.22, 3.30) | 5.81E-03 |
| TMEM95 | 17p13.3 | 7199221-7201262 | 1.25(1.06, 1.46) | 7.24E-03 |
| PHF2 | 9q22.31 | 95378730-95481690 | 0.81(0.69, 0.94) | 7.30E-03 |
| RHOJ | 14q23.2 | 62740898-62828312 | 0.78(0.65, 0.94) | 7.60E-03 |
| SLC1A7 | 1p32.3 | 53325443-53380837 | 1.24(1.06, 1.46) | 8.71E-03 |
| GRIK2 | 6q16.3 | 101953626-102624651 | 0.78(0.65, 0.94) | 8.74E-03 |
| DACH1 | 13q21.33 | 70910099-71339331 | 0.82(0.70, 0.95) | 1.08E-02 |
| AFG1L | 6q21 | 108722951-108953897 | 0.77(0.63, 0.94) | 1.13E-02 |
| SEMA3E | 7q21.11 | 82831158-83116260 | 0.82(0.70, 0.96) | 1.18E-02 |
| TOX | 8q12.1 | 59880531-60194321 | 1.51(1.09, 2.09) | 1.29E-02 |
| EXT1 | 8q24.11 | 118880783-119193239 | 0.62(0.42, 0.91) | 1.41E-02 |
| TNFSF13B | 13q33.3 | 107719978-107757366 | 0.82(0.69, 0.96) | 1.48E-02 |
| RPH3AL | 17p13.3 | 62294-202576 | 1.20(1.03, 1.40) | 2.13E-02 |
| VWA1 | 1p36.33 | 1360772-1366009 | 1.71(1.07, 2.73) | 2.41E-02 |
| CYHR1 | 8q24.3 | 145646123-145661287 | 1.62(1.06, 2.47) | 2.43E-02 |
| SCN3A | 2q24.3 | 165652276-165768823 | 0.82(0.69, 0.98) | 2.62E-02 |
| TPK1 | 7q35 | 143779967-144164079 | 0.82(0.68, 0.98) | 2.75E-02 |
| NBEAL1 | 2q33.2 | 203708718-203790962 | 1.23(1.02, 1.47) | 2.78E-02 |
| ZNF79 | 9q33.3 | 129226474-129247472 | 1.24(1.02, 1.51) | 2.81E-02 |
| ASB1 | 2q37.3 | 239000365-239025630 | 1.21(1.02, 1.44) | 2.90E-02 |
| TSPAN8 | 12q21.1 | 69805144-69838046 | 0.84(0.72, 0.98) | 2.98E-02 |
| RBM25 | 14q24.2 | 72595021-72657745 | 0.79(0.64, 0.98) | 2.99E-02 |

| | | | | |
|---|---|---|---|---|
| IMPG1 | 6q14.1 | 76687782-76839055 | 0.84(0.72, 0.98) | 3.01E-02 |
| FAM110D | 1p36.11 | 26358157-26363040 | 0.84(0.71, 0.98) | 3.19E-02 |
| GYPC | 2q14.3 | 127130154-127170716 | 1.18(1.01, 1.38) | 3.20E-02 |
| RPL27AP6 | 6q25.2 | 154000331-154002595 | 0.84(0.71, 0.99) | 3.24E-02 |
| LINC00638 | 14q32.33 | 104358583-104361100 | 1.50(1.03, 2.18) | 3.29E-02 |
| TRIQK | 8q22.1 | 93964938-94047548 | 0.81(0.66, 0.99) | 3.53E-02 |
| PGAM5 | 12q24.33 | 131797509-131805825 | 1.62(1.03, 2.54) | 3.75E-02 |
| ACSL5 | 10q25.2 | 114125946-114178128 | 0.85(0.73, 0.99) | 3.83E-02 |
| F7 | 13q34 | 112808106-112822996 | 1.57(1.02, 2.42) | 4.00E-02 |
| DAD1 | 14q11.2 | 22103647-22127983 | 0.85(0.73, 0.99) | 4.27E-02 |
| MEP1B | 18q12.1 | 28023985-28054364 | 0.80(0.65, 0.99) | 4.34E-02 |
| PTPRN2 | 7q36.3 | 157024516-158073179 | 1.17(1.00, 1.37) | 4.37E-02 |
| ADAM6 | 14q32.33 | 105506864-105509403 | 1.17(1.00, 1.37) | 4.40E-02 |
| TCERG1L | 10q26.3 | 132780645-132999974 | 0.85(0.73, 1.00) | 4.53E-02 |
| SLITRK5 | 13q31.2 | 87122871-87129871 | 0.85(0.72, 1.00) | 4.61E-02 |
| SPHKAP | 2q36.3 | 228552919-228754586 | 0.86(0.73, 1.00) | 4.64E-02 |
| FBXO21 | 12q24.22 | 116065968-116112683 | 0.85(0.73, 1.00) | 4.76E-02 |
| RIN2 | 20p11.23 | 19818210-19931100 | 0.82(0.67, 1.00) | 4.82E-02 |
| FAM155A | 13q33.3 | 106618880-107317084 | 0.84(0.70, 1.00) | 4.94E-02 |

**Supplementary Table S2. Significantly associated germline CNVRs with melanoma risk detected by Model 2.** CNV calls were first generated by using modSaRa, and CNVRs were then defined by using RO method by using CNVRuler. OSCAA was applied to each CNVR adjusting age and gender effect. NCBI build 36 (hg18) was used to find overlapped genes for CNVRs with significant CNV-disease association (p value < 0.05). 26 significant genes were shown in the table. RO: reciprocal overlap; CNVR: CNV region.

| Gene | Chr | Coordinates | OR (95% CI) | P value |
|---|---|---|---|---|
| HSF1 | 8q24.3 | 145486078-145509193 | 4.05(1.89, 8.67) | 0.00E+00 |
| DGAT1 | 8q24.3 | 145510762-145521375 | 4.61(2.06, 10.33) | 0.00E+00 |
| PTPN11 | 12q24.13 | 111340919-111432100 | 15.61(3.87, 62.90) | 0.00E+00 |
| MLPH | 2q37.3 | 238060617-238128700 | 6.24(2.53, 15.37) | 0.00E+00 |
| ARID1B | 6q25.3 | 157140778-157572094 | 2.87(1.60, 5.15) | 0.00E+00 |
| ANKLE2 | 12q24.33 | 131812327-131848524 | 2.54(1.46, 4.42) | 1.00E-03 |
| C1ORF159 | 1p36.33 | 1007061-1041599 | 2.54(1.45, 4.47) | 1.00E-03 |
| TECPR1 | 7q21.3 | 97683983-97719404 | 2.35(1.41, 3.93) | 1.00E-03 |
| HACE1 | 6q21 | 105282661-105414487 | 0.66(0.49, 0.88) | 4.00E-03 |
| ERBB4 | 2q34 | 211948687-213111597 | 0.74(0.60, 0.91) | 4.00E-03 |
| TTLL10 | 1p36.33 | 1099146-1123176 | 2.17(1.29, 3.65) | 4.00E-03 |
| TCF15 | 20p13 | 532637-538910 | 2.11(1.25, 3.59) | 5.00E-03 |
| CDC34 | 19p13.3 | 482733-493087 | 2.01(1.22, 3.30) | 6.00E-03 |
| PWWP2B | 10q26.3 | 134060692-134081348 | 1.80(1.16, 2.79) | 9.00E-03 |
| TMCC1 | 3q21.3 | 130849325-131095093 | 0.63(0.44, 0.90) | 1.10E-02 |
| EXT1 | 8q24.11 | 118915232-118952404 | 0.62(0.42, 0.91) | 1.40E-02 |
| TNFSF13B | 13q33.3 | 107719978-107757366 | 0.81(0.69, 0.96) | 1.40E-02 |
| MDGA2 | 14q21.3 | 46378578-47213738 | 0.73(0.57, 0.94) | 1.50E-02 |
| C2CD4C | 19p13.3 | 356445-360147 | 1.87(1.12, 3.13) | 1.70E-02 |
| CRIP2 | 14q32.33 | 105012176-105017545 | 1.40(1.05, 1.85) | 2.10E-02 |
| TNFRSF18 | 1p36.33 | 1128751-1131952 | 1.73(1.09, 2.75) | 2.10E-02 |
| NCOA7 | 6q22.32 | 126153694-126293959 | 0.73(0.55, 0.96) | 2.30E-02 |
| PGAM5 | 12q24.33 | 131797509-131805825 | 1.62(1.03, 2.54) | 3.70E-02 |
| GRM8 | 7q31.33 | 125865888-126670805 | 0.84(0.71, 0.99) | 3.80E-02 |
| TPK1 | 7q35 | 143779967-144164079 | 0.83(0.69, 1.00) | 4.90E-02 |
| PRORP | 12q13.2 | 34661509-34816579 | 1.19(1.00, 1.41) | 4.90E-02 |
