## Supplementary material for "OSCAA: A Two-Dimensional Gaussian Mixture Model for Copy Number Variation Association Analysis": Figures and Tables

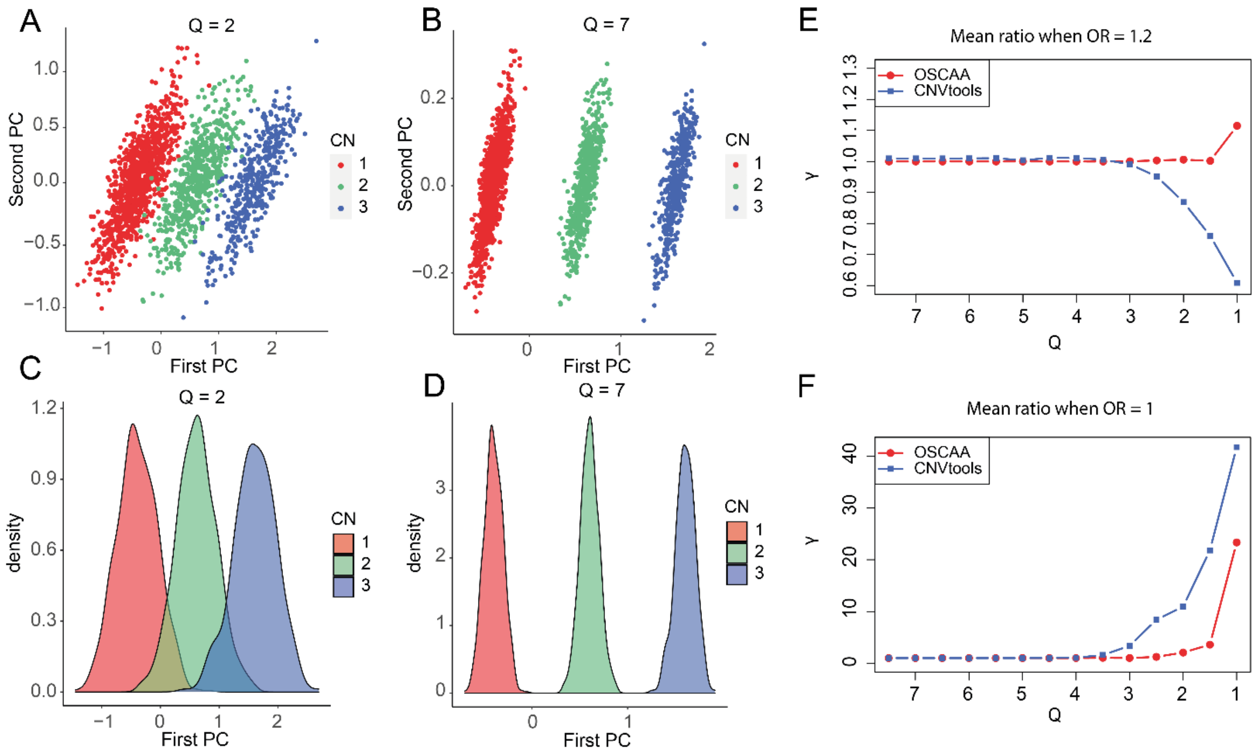


**Fig. 1. Performance evaluation of OSCAA via simulation studies in comparison to one-stage method.** Assume there was a correlation between the first and second PCs (correlation coefficient $\rho$ = 0.8), signals for 2,000 samples were simulated for 3 different CNV states: deletions, normal, and duplications from bivariate normal distributions with parameters derived based on $Q$. The CNV proportion varied among 50%, 30%, and 20%. $Q$ denoted the signal-to-noise ratio ranging from 1 to 7.5. A binary disease status was simulated; OSCAA was fitted using the first two PCs while CNVtools was fitted by only using the first PC. Distribution of CNV signals were displayed for $Q$ = 2 (A, C) and 7 (B, D), respectively. Mean ratio ($\gamma$) was used to evaluate the performance of each method under various $Q$ values when OR = 1.2 (E) and OR = 1 (F). $\gamma$: mean ratio between estimated $\chi^2$ and true $\chi^2$ across 100 simulations. OR: Odds Ratio.

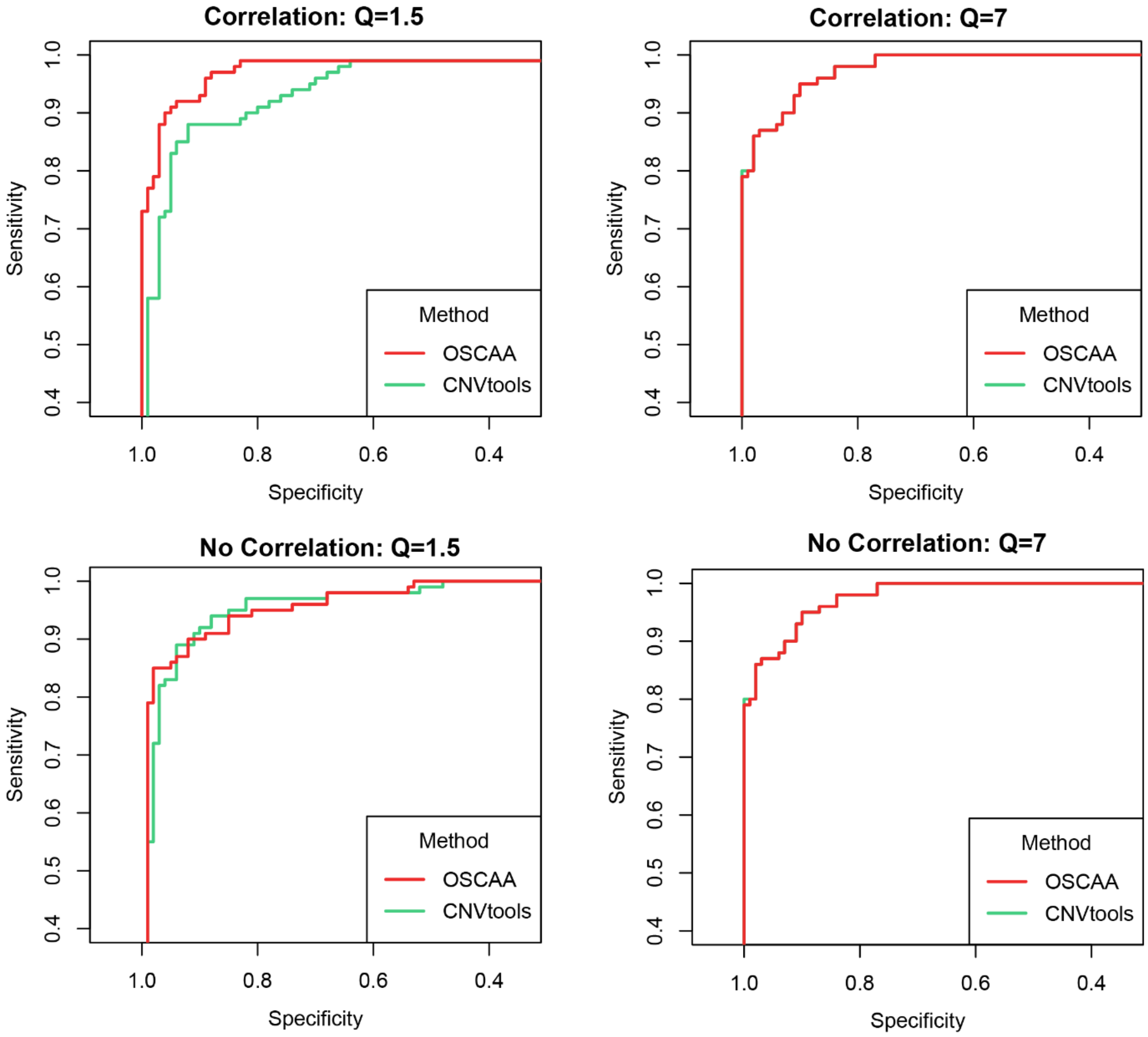


**Fig. 2. Performance evaluation of OSCAA via simulation studies**. Simulation studies were conducted for various combination of $Q$ and $\rho$, where $Q$ denoted the signal-to-noise ratio ranging from 1 to 7.5 and $\rho$ was the correlation between two PCs with $\rho$ = 0.8 for a positive correlation between while $\rho$ = 0 when there was no correlation. ROC curve was used to evaluate the performance of OSCAA and CNVtools in four scenarios: there was a correlation between PCs when $Q$ = 1.5 (A) and $Q$ = 7 (B); there was no correlation between PCs when $Q$ = 1.5 (C) and $Q$ = 7 (D).

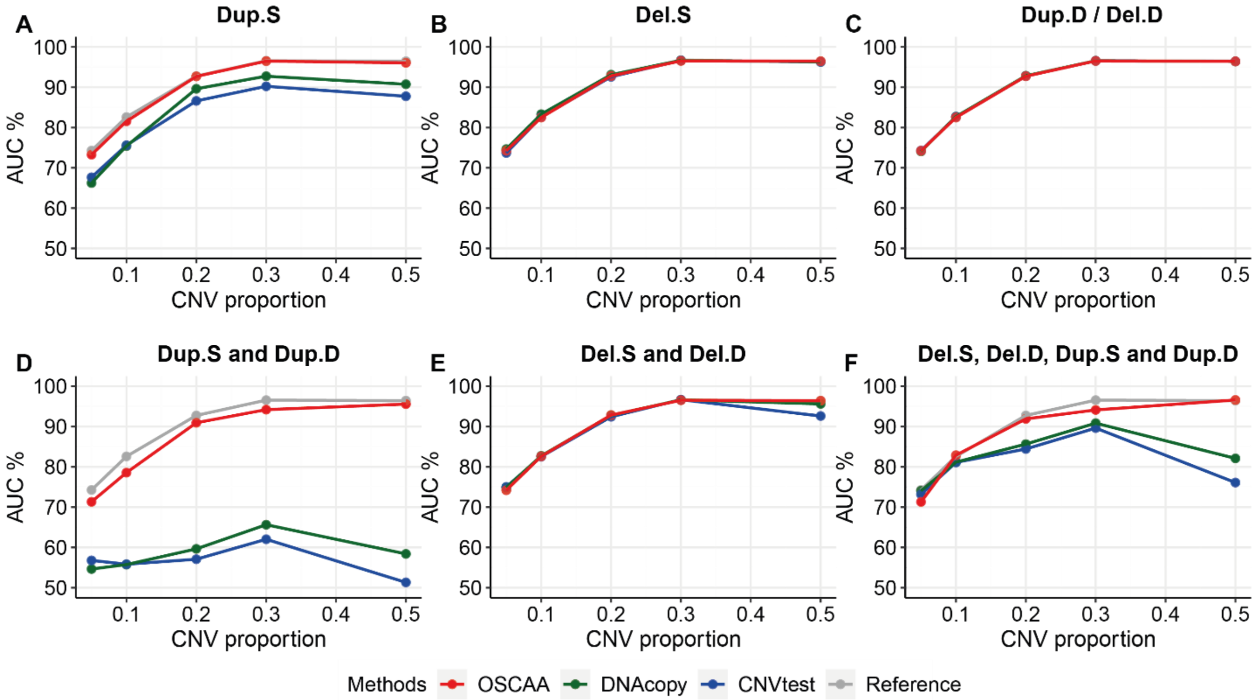


**Fig. 3. Performance evaluation of OSCAA with two-stage methods in short CNVs via simulation studies with $OR_{h1}$ = 1.5.** Signals of 120 CNVRs of short CNV length (5~20 probes) were simulated for varied copy number states including Dup.S (A), Del.S (B), Del.D/ Dup.D (C), mixture of 2 duplication states (D: Dup.S and Dup.D), mixture of 2 deletion states (E: Del.D and Del.S); and mixture of 4 CNV states (F: Del.D, Del.S, Dup.S and Dup.D). The CNV proportion ranged from 0.05 to 0.5. In each scenario, a binary disease status was simulated for each sample with OR = 1.5 or OR = 1. The reference line (grey line) represented the actual AUCs with each CNV correctly detected. Del.D: deletion of double copies; Del.S: deletion of single copy; Dup.S: duplication of single copy; Dup.D: duplication of double copies; OR: Odds Ratio; SD: standard deviation; CNVR: CNV region; AUC: area under curve.

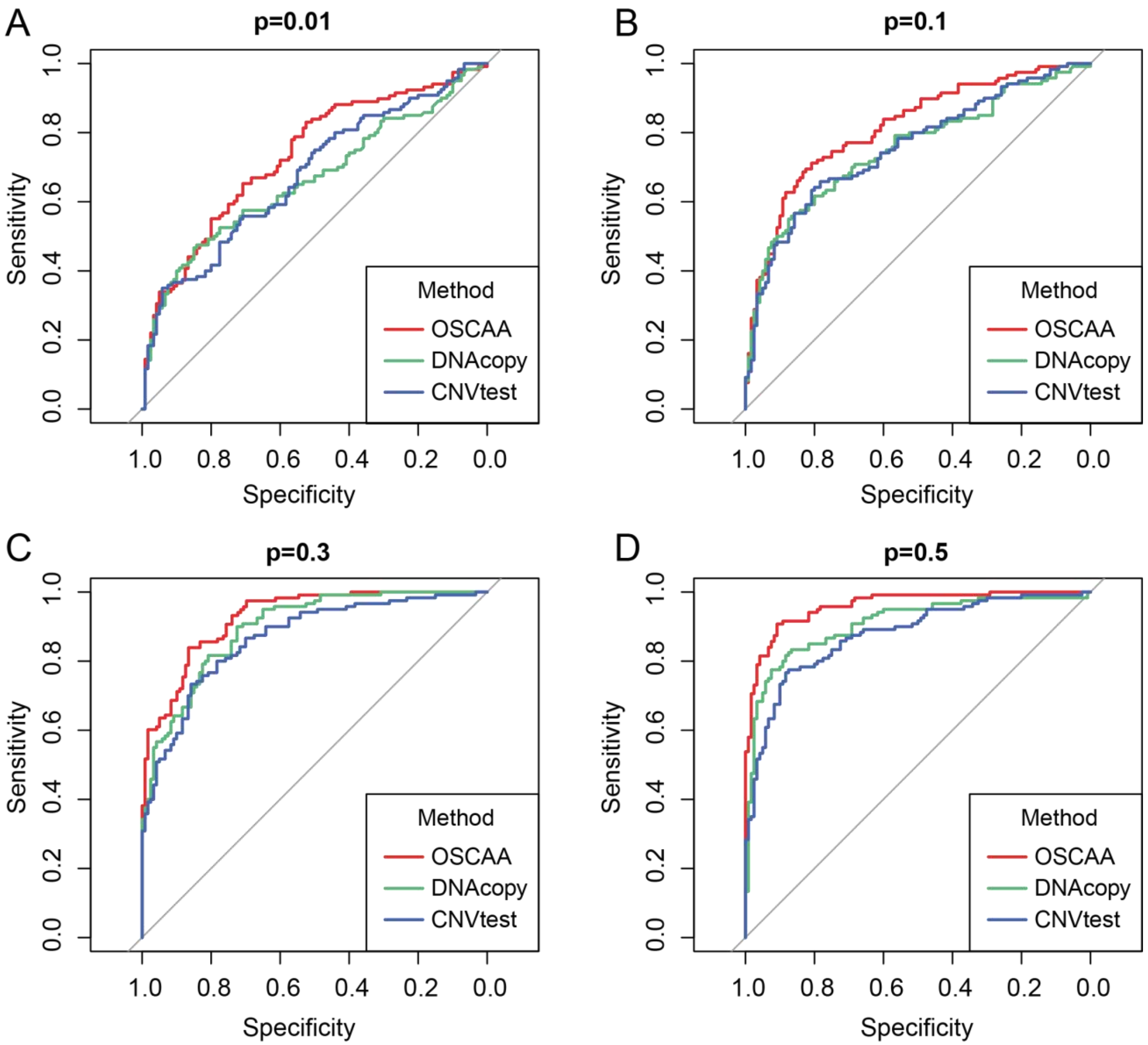


**Fig. 4. Performance evaluation of OSCAA with two-stage methods, DNAcopy and CNVtest in short Dup.S with $OR_{h1}$ = 1.5.** Signals of 120 Dup.S of short CNV length (5~20 probes) were simulated with CNV proportion ranging from 0.05 to 0.5. A binary disease status was then simulated for each sample from a binomial distribution with OR = 1.5 or OR = 1. Dup.S: duplication of single copy; OR: Odds Ratio; ROC: receiver operating characteristic.

**Table 1. Top significantly associated germline CNVRs with melanoma risk detected by OSCAA using Model 1.** CNV calls were first generated by using modSaRa, and CNVRs were then defined by using RO method by using CNVRuler. OSCAA was applied to each CNVR adjusting age and gender effect, assuming same risk effect for deletions and duplications. NCBI build 36 (hg18) was used to find overlapped genes for CNVRs with significant CNV-disease association (p value < 0.05). Top 28 significant genes were shown in the table. RO: reciprocal overlap; CNVR: CNV region.

| Gene | Chr | Coordinates | OR (95% CI) | P value |
|---|---|---|---|---|
| MXRA8 | 1p36.33 | 1275872-1285186 | 1.47(1.24, 1.76) | 1.54E-05 |
| GPR35 | 2q37.3 | 241214309-241218002 | 2.72(1.57, 4.73) | 3.80E-04 |
| TXNDC2 | 18p11.22 | 9797167-9806111 | 0.45(0.29, 0.70) | 4.59E-04 |
| MACROD2 | 20p12.1 | 15249783-15251701 | 0.76(0.65, 0.89) | 5.66E-04 |
| C1ORF159 | 1p36.33 | 1064387-1069061 | 2.61(1.49, 4.56) | 7.71E-04 |
| ARFGEF1 | 8q13.2 | 68289797-68327429 | 0.63(0.48, 0.83) | 7.76E-04 |
| ARID1B | 6q25.3 | 157487691-157510801 | 2.39(1.41, 4.05) | 1.23E-03 |
| OR7C2 | 19p13.12 | 14907383-14908117 | 1.29(1.10, 1.50) | 1.29E-03 |
| TECPR1 | 7q21.3 | 97705969-97711427 | 1.26(1.08, 1.48) | 4.18E-03 |
| PWWP2B | 10q26.3 | 134050145-134064927 | 1.27(1.08, 1.51) | 4.50E-03 |
| OR4N2 | 14q11.2 | 19365350-19397391 | 2.52(1.32, 4.82) | 5.08E-03 |
| TCF15 | 20p13 | 531587-541641 | 2.12(1.25, 3.59) | 5.46E-03 |
| NCS1 | 9q34.11 | 132066424-132219132 | 1.28(1.08, 1.53) | 5.68E-03 |
| CDC34 | 19p13.3 | 460897-487878 | 2.01(1.22, 3.30) | 5.81E-03 |
| TMEM95 | 17p13.3 | 7205517-7206463 | 1.25(1.06, 1.46) | 7.24E-03 |
| PHF2 | 9q22.31 | 95540187-95541151 | 0.81(0.69, 0.94) | 7.30E-03 |
| RHOJ | 14q23.2 | 62793874-62794338 | 0.78(0.65, 0.94) | 7.60E-03 |
| SLC1A7 | 1p32.3 | 53366758-53368090 | 1.24(1.06, 1.46) | 8.71E-03 |
| GRIK2 | 6q16.3 | 102041883-102044666 | 0.78(0.65, 0.94) | 8.74E-03 |
| DACH1 | 13q21.33 | 71364644-71394627 | 0.82(0.70, 0.95) | 1.08E-02 |
| AFG1L | 6q21 | 108719169-108721409 | 0.77(0.63, 0.94) | 1.13E-02 |
| SEMA3E | 7q21.11 | 82888668-82889067 | 0.82(0.70, 0.96) | 1.18E-02 |
| TOX | 8q12.1 | 60289264-60298951 | 1.51(1.09, 2.09) | 1.29E-02 |
| EXT1 | 8q24.11 | 118915232-118952404 | 0.62(0.42, 0.91) | 1.41E-02 |
| TNFSF13B | 13q33.3 | 107745089-107750000 | 0.82(0.69, 0.96) | 1.48E-02 |
| RPH3AL | 17p13.3 | 193933-196701 | 1.20(1.03, 1.40) | 2.13E-02 |
| VWA1 | 1p36.33 | 1346413-1380091 | 1.71(1.07, 2.73) | 2.41E-02 |
| CYHR1 | 8q24.3 | 145659045-145662349 | 1.62(1.06, 2.47) | 2.43E-02 |
